# Genomic and Phenotypic Characterization of Mupirocin Resistant *Staphylococcus aureus* Clinical Isolates

**DOI:** 10.1101/2025.04.16.648996

**Authors:** Ariana M. Virgillio, Emily A. Felton, Jessica K. Jackson, Sarah J. Kennedy, Deanna N. Becker, Amorce Lima, Kimberly Atrubin, Eleonora Cella, Taj Azarian, Suzane Silbert, Lindsey N. Shaw, Kami Kim

## Abstract

**Background:** Colonization with *Staphylococcus aureus* is a risk factor for subsequent infection. Decolonization with the topical antibiotic mupirocin is effective and reduces the risk of subsequent *S. aureus* infection for both methicillin-sensitive (MSSA) and methicillin-resistant (MRSA) strains but may select for mupirocin-resistant isolates.

**Methods:** We characterized oxacillin and mupirocin susceptibility amongst 384 *S. aureus* strains isolated from clinical samples isolated 2017-2023 in Tampa, Florida, spanning strains collected before and after the onset of the COVID-19 pandemic. Whole genome sequencing of bacterial isolates was conducted in parallel and correlated with drug susceptibility profiles.

**Results:** Mupirocin resistance (MupR) was nearly exclusively present in MRSA strains (103/106 97.1% of MupR; 103/299 34.4% of MRSA). Although our hospital protocol for decolonization shifted to povidone iodine in the Post-COVID period, the overall prevalence of MupR did not change in Pre-COVID and Post-COVID samples (28.9% vs 26%). Genotype correlated with antibiotic susceptibility with low level MupR (MupLR), linked to mutations in *ileS* and high level MupR (MupHR), linked to the presence of *mupA*. Genome analysis revealed that most MupR strains fell into three sequence types (ST) falling into two major clonal complexes (CC): CC8 ST8 (including Community-Associated MRSA strains USA300 and USA500), CC5 ST5 (associated with Healthcare-Associated MRSA such as USA100), and CC5 ST3390. ST3390 isolates had the highest prevalence of MupR (30/36 83%; MupHR 20/36 55.6%; MupLR 10/36 27.8%).

**Conclusions:** Mupirocin resistance was prevalent in our hospital MRSA strains. We also found evidence for emergence and persistence of ST3390 MRSA-MupR strains in Florida.

**Key points:**

- In a survey of clinical isolates in Florida, 34.4% of MRSA strains were mupirocin resistant.
- Mupirocin resistance correlated with mutations in *ileS* or carriage of *mupA*.
- We found evidence for emergence of MRSA mupirocin-resistant strains that were sequence type ST3390.

## Introduction

Infections with *Staphylococcus aureus* are major causes of skin and soft tissue infections, as well as invasive infections such as endocarditis, pneumonia, and osteomyelitis. Colonization of skin and mucosal surfaces with *S. aureus* is a risk factor for subsequent clinical infections [1]. Decolonization is an effective method for reducing risk of subsequent *S.aureus* infections, including those caused by methicillin resistant *S. aureus* (MRSA) [2]. Universal decolonization with mupirocin and chlorhexidine body washes has been shown in randomized control trials as being more effective than targeted decolonization [3]. Decolonization of *S. aureus* prevents post-surgical infections, ventilator associated pneumonias and blood stream infections [4].

Mupirocin inhibits bacterial protein synthesis by competitively binding to the isoleucine specific site of the isoleucyl-tRNA synthetase preventing the conversion of isoleucine and tRNA to isoleucyl-tRNA. As this is an essential process, this inhibition leads to bacterial cell death. There have been reports of increasing prevalence of mupirocin resistance, particularly amongst MRSA strains, hypothesized due to increased usage of decolonization protocols designed to prevent MRSA infections [5].

Mupirocin resistance (MupR) can be categorized as mupirocin susceptible (MupS, ≤4 µg/mL), low-level resistant (MupLR, 8-64 µg/mL), and high-level resistant (MupHR, ≥ 512 µg/mL) [6] [7] [8] [9] [10] [11]. Isolates demonstrating MupLR typically have a chromosomal mutation altering the Rossman fold of the native isoleucyl-tRNA synthetase (IleS) protein sequence [12]. The most prevalent mutations are V588F and V631F, which result in bulkier residue within the enzymatic binding site, blocking association with mupirocin. The major determining factor behind high-level resistance is the gene *mupA*, although exceptions have been noted, including *mupA*-negative MupHR-isolates that possess a different gene, *mupB*, which is 65% similar to *mupA* [13]. Whilst mupirocin binds discriminately to the substrate binding sites of bacterial isoleucyl-tRNA synthetases, it has low affinity for eukaryotic isoleucyl-tRNA synthetases. Both MupA and MupB have altered sequences that are closer to that of eukaryotes, whilst still functional in prokaryotes. Importantly, both genes can be found on multi-resistant plasmids, facilitating the spread of resistance.

In this study we apply a combination of Whole Genome Sequencing (WGS) and phenotypic characterization to investigate changes in the molecular epidemiology of *S. aureus* mupirocin resistance within a large metropolitan hospital in Florida. Our study period includes isolates before and after the COVID-19 pandemic, which coincided with a switch in hospital protocols from mupirocin to povidone iodine for decolonization.

## Methods

### SA Strain Sampling

A random convenience sample of clinical *S. aureus* isolates was collected from discarded residual clinical samples at Tampa General Hospital (TGH) between 2017-2023. TGH is a 1040 bed not-for-profit academic medical center serving a population exceeding six million across western Florida. The TGH research lab archives more MRSA than MSSA isolates and randomly archives samples from different body sites.

### Patient Consent Statement

Collection of samples for this study was reviewed by our local IRB and was deemed not human subjects research and did not require patient consent.

### MIC Testing

Samples were struck to TSA for isolation overnight and transferred to TSB for overnight growth at 37°C in a shaking incubator. MIC testing was performed using a top agar method. Clinical susceptibility data were verified using E-tests MupR was designated as sensitive (MIC <4 µg/mL), low-level or intermediate resistant (MupLR, MIC = 8-256 µg/mL), or high level resistant (MupHR, MIC >512 µg/mL). Bacteria were suspended to 0.5 McFarland in Mueller Hinton II agar for mupirocin testing and Muller Hinton II Agar + 2% NaCl for oxacillin testing. Inoculated agar (10mL) was poured over premade uninoculated plates with their corresponding agar. Test strips from Liofilchem were placed on dried agar corresponding to the antibiotic being tested. Muller Hinton II agar with mupirocin test strips were incubated inverted at 37°C for 18 hours. Muller Hinton II +2% NaCl agar with oxacillin test strips were incubated inverted at 37°C for 24 hours. Mupirocin plates were read with the guidance of “MIC Test Strip Photographic Guide” with interpretation as in current literature. Oxacillin plates were read using CLSI breakpoints provided by Liofilchem. Analysis for statistical differences between groups was performed with SPSS using χ² for significance (p=0.05).

### Whole Genome Sequencing (WGS)

Overnight cultures were grown in TSB at 37⁰C with shaking. Genomic DNA was collected using the DNeasy Blood and Tissue kit (Qiagen) following the gram-positive specific pretreatment method, with the addition of 20mg/ml lysostaphin. For 80 strains (Run 1), quality and quantity was assessed using a Qubit 4 Fluorometer, and sequencing occurred on an Illumina MiSeq instrument with a Nextera Flex library preparation kit and V3 flow cell chemistry. For the remaining 304 strains, quality and quantity was assessed using a Quant-it fluorescent assay kit (Invitrogen) and a Biotek Citation5 microplate reader. Libraries were prepared using a Hackflex [14] adapted version of the Illumina DNA-seq library preparation kit. Sequencing of 70 of the Pre-COVID samples (Run 2) was performed using a Nextseq550 Mid-Output v2.5 Kit (300 cycles) and 234 Post-COVID samples were sequenced using a Nextseq2000 with P1 flow cell chemistry (Run 3). Runs 2 and 3 were performed at the USF Genomics Center Core facilities. Strains are listed in Supplementary Table 1.

For long read sequencing of TGH462, DNA was extracted using the QIAGEN genomic tip 100/G and genomic DNA buffer kits. Genomic DNA integrity and concentration were accessed via qubit 4 (Invitrogen) and Genomic DNA Tapestation (Agilent) QC checks. Samples were then treated with the short fragment eliminator (Nanopore) and prepared with the Rapid Barcoding library prep kit (SQK-RBK114.24). Libraries were sequenced on an R10.4.1 flowcell using a MinION Mk1B with default settings of 72 hour run length and 200 bp minimum read length.

### Genomic assembly and molecular characterization

The raw reads for all samples were trimmed via Trimmomatic v0.39 [15] with SLIDINGWINDOW:10:20 MINLEN:31 TRAILING:2 settings [16]. Trimming quality checks were performed with FastQC v0.12.0. The reads were then *de novo* assembled with SPAdes v3.15.5 using default settings [17], and assemblies were checked for quality via Quast v5.2.0 [18]. Genomes were then sequence-typed with pyMLST [19], annotated with prokka, and antimicrobial resistance (AMR) genes identified using AMRFinderPlus v3.10.45 [20]. The presence of DNA and translated Protein sequences for AMR genes of interest were confirmed and aligned using BLASTncbi-BLAST v2.13.0+ [21]. The staphtyper workflow as part of Bactopia v3.0.1 was used to further characterize isolates based on Agr group [22], SCCmec type [23], and Spa type [24].

### Phylogenetic analysis

To explore relatedness of the cluster of ST8-t008 that harbor the novel R112Q *ileS* mutation, Snippy was used to generate an initial core genome single-nucleotide polymorphism alignment. Following this, Gubbins was used to create a recombination masked alignment. From this alignment RAxML was used to infer a maximum-likelihood mid-point rooted tree. The subsequent tree was visualized and annotated using iTOL webserver.

## Results

### Mupirocin Resistance (MupR) is Common Amongst Tampa MRSA isolates

To explore the prevalence of MupR in our hospital system, we curated and characterized a collection of 384 *S. aureus* strains. The collection was isolated between February 2017 and September 2023, and comprised of 299 MRSA isolates (77.9%) and 85 MSSA strains (22.1%) (**Figure 1**). Many infection control practices changed during the COVID-19 pandemic, and there have been reports of increased *S. aureus* incidence during the pandemic [25, 26] [27]. Therefore, we designated the collection as being collected Pre-COVID (**Table 1**; 2017-2020, PreC = 211 strains, 54.9%) and Post-COVID (2021-2023, PostC = 173 strains, 45.1%). There were 161 Blood samples making up 41.9% of the sample set (**Table 1)**. Non-Blood strains (223 strains, 58.1%) were isolated from infections of: bone (4 samples, 1%), eye (12 samples, 3.1%), fluid (29 samples, 7.6%), respiratory (58 samples, 15.1%), urine (37 samples, 9.6%), and wound (83 samples, 21.6%) (**Table 1**).

**Figure 1.**
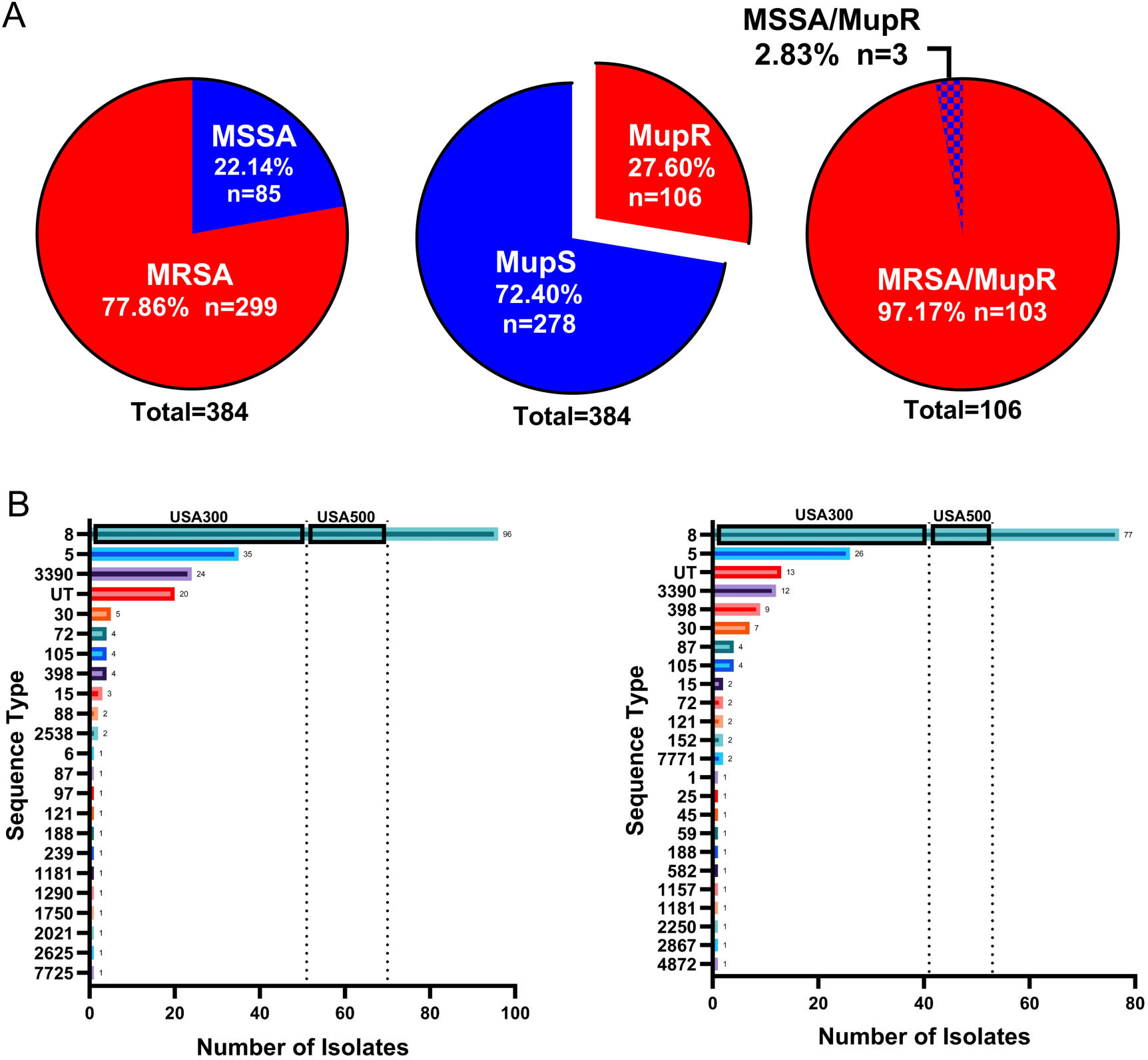
Mupirocin Resistance and Sequence Typing of Tampa *S. aureus* isolates. A. 384 *S.aureus* clinical strains from 2017-2023 were characterized. Oxacillin sensitivity (MSSA, n=85), mupirocin sensitivity (MupS, n=278) and the distribution of MRSA and MSSA strains amongst mupirocin resistant (MupR n=106) strains is shown. B. Results of classification of strains into sequence types (ST) are shown with numbers of Pre-COVID isolates (2017-2020) and Post-COVID isolates (2021-2023) within each ST. Numbers of USA300 and USA500 within ST8 are shown. ST5 includes USA100 strains. “UT” signifies strains that were untypeable using the methods deployed herein. ST was determined by analysis of whole genome sequence of each strain.

**Table 1.** *S. aureus* strains used in this study. Numbers collected Pre-COVID (2017-2020) and Post-COVID (2021-2023) are indicated, as are the numbers of strains in each group with oxacillin resistance and mupirocin resistance as defined in the methods. Sites of collection of the strains are also indicated. All strains underwent whole genome sequencing (details in supplementary table 2)

|  | *Strains* | *Ox R(MRSA)* | *Mup R* |
| --- | --- | --- | --- |
| ***Pre-COVID*** | 211 | 164 | 61 |
| ***Post-COVID*** | 173 | 135 | 45 |
| ***Blood*** | 161 | 100 | 35 |
| ***Non-Blood*** | 223 | 199 | 71 |
| ***Total*** | 384 | 299 | 106 |

**Strain Source**
|  | ***PreC*** |  |  | ***PostC*** |  |  | ***Total*** |
| --- | --- | --- | --- | --- | --- | --- | --- |
|  | MupHR | MupLR | MupS | MupHR | MupLR | MupS | No (%) |
| *Total #* | 44 | 17 | 150 | 28 | 17 | 128 | 384 (100) |
| *Blood* | 12 | 5 | 65 | 9 | 9 | 61 | 161 (41.9) |
| *Non-Blood* | 32 | 12 | 85 | 19 | 8 | 67 | 223 (58) |
| *Bone* | 1 | 0 | 2 | 0 | 0 | 1 | 4 (1) |
| *Eye* | 1 | 0 | 1 | 6 | 0 | 4 | 12 (3.1) |
| *Fluid* | 3 | 1 | 12 | 1 | 0 | 12 | 29 (7.6) |
| *Respiratory* | 13 | 5 | 21 | 2 | 3 | 14 | 58 (15.1) |
| *Urine* | 5 | 1 | 12 | 4 | 2 | 13 | 37 (9.6) |
| *Wound* | 9 | 5 | 37 | 6 | 3 | 23 | 83 (21.6) |

The Pre-COVID and Post-COVID periods coincided with switch from a mupirocin-based decolonization protocol to povidone-based protocol in January 2020. Our hospital had found >40% prevalence of mupirocin resistance in 2019 in amongst MRSA isolates from surveillance nasal swabs and a random sampling of MRSA blood cultures (46% of MRSA nares n=127; 43% of MRSA blood cultures n=46). The first case of COVID-19 in the Tampa Bay area was March 1, 2020 with Post-COVID cases collected more than a year after change in decolonization protocol. We determined the prevalence of mupirocin resistance amongst both MRSA and MSSA isolates and whether specific genetic or phenotypic features correlated with resistance over time.

All strains were screened for oxacillin and mupirocin resistance using Etest strips. From this, we determined that 278 of 384 strains (72.4%; **Figure 1A**, **Table 1**) were susceptible to mupirocin. Of the 106 (27.6%) resistant strains, 34 (8.9%) had low-level resistance, whilst 72 (18.8%) had high level resistance.

### Mupirocin Resistance Prevalence was Stable

We observed significant correlation between oxacillin resistance and mupirocin resistance (**Supplemental Table 2 and 3**). Most MupR strains were MRSA (**Figure 1A**). From Pre-COVID samples, 2.1% of MSSA samples were also MupR, whilst 36.6% of MRSA samples were MupR (p<0.001). 30.7% of MupS samples were oxacillin susceptible, whilst 98.4% of MupR samples were oxacillin resistant. (p<0.001). Post-COVID strain susceptibilities were similar. For example, 5.3% of MSSA samples were MupR and 31.9% of MRSA samples were MupR (p<0.001).

When comparing levels of MupR between the Pre- and Post-COVID time points, there was no significant difference between the two strain sets (p=0.460). In the Pre-COVID population, 71.1% of strains were susceptible, whilst 28.9% displayed resistance (MupLR = 8.1%, MupHR = 20.9%). In the Post-COVID strains, 74.0% were susceptible, whilst 26% displayed resistance (MupLR = 9.8%, MupHR = 16.2%).

### Pre-COVID Non-Blood Strains Had Higher Levels of MupR

We found a significant difference for strains isolated from blood versus non-blood. 21.7% of blood isolates displayed some level of MupR, compared with 31.8% of non-blood samples (p=0.029; **Supplemental Table 4 and 5**). This was driven by the Pre-COVID population, where 20.7% of blood samples displayed resistant versus 34.1% of non-blood samples (p=0.037), whilst Post-COVID had no significant correlation between isolate site and resistance (22.8% of blood samples were resistant versus 28.7% of non-blood samples, p=0.375). When the non-blood strains were parsed more deeply, 46.2% of the Pre-COVID respiratory isolates were MupR (p=0.016), which accounts for 29.5% of all Pre-COVID MupR strains. Amongst our Post-COVID cohort, 26.3% respiratory strains were MupR, making up 11.1% of MupR samples in the Post-COVID era (p=0.651).

### Clonal Clusters and Sequence Types Associated with MupR

All strains from our collection were subject to WGS and initially parsed into CC and then ST (**Figure 1B**). Strains that share six of the seven conserved sequences used for multilocus sequence typing (MLST) are grouped into clonal complexes (CC). MupR was most common in three sequence types (**Figure 2**). CC8 ST8 was the most abundant, with 173 strains (PreC = 96, 45.5% vs. PostC = 77, 44.5%), followed by 61 CC5 ST5 strains (PreC = 35, 16.6% vs. PostC = 26, 15%) and 36 CC5 ST3390 strains (PreC = 24, 11.4% vs. PostC = 12, 7.5%). CC8 ST8 encompass USA300 and USA500, the most common CA-MRSA in the US. There were 92 USA300 (ST8 t008) isolates (PreC = 51, 55.4% vs. PostC = 41, 44.6%), and 31 USA500 (ST8 t064) strains (PreC = 19, 61.3% vs. PostC = 12, 38.7%). CC5 ST5 includes USA100, the CC/ST type classically associated with HA-MRSA) in the US [28]. ST5, ST3390 and ST105 are closely related strains that are members of CC5.

**Figure 2.**
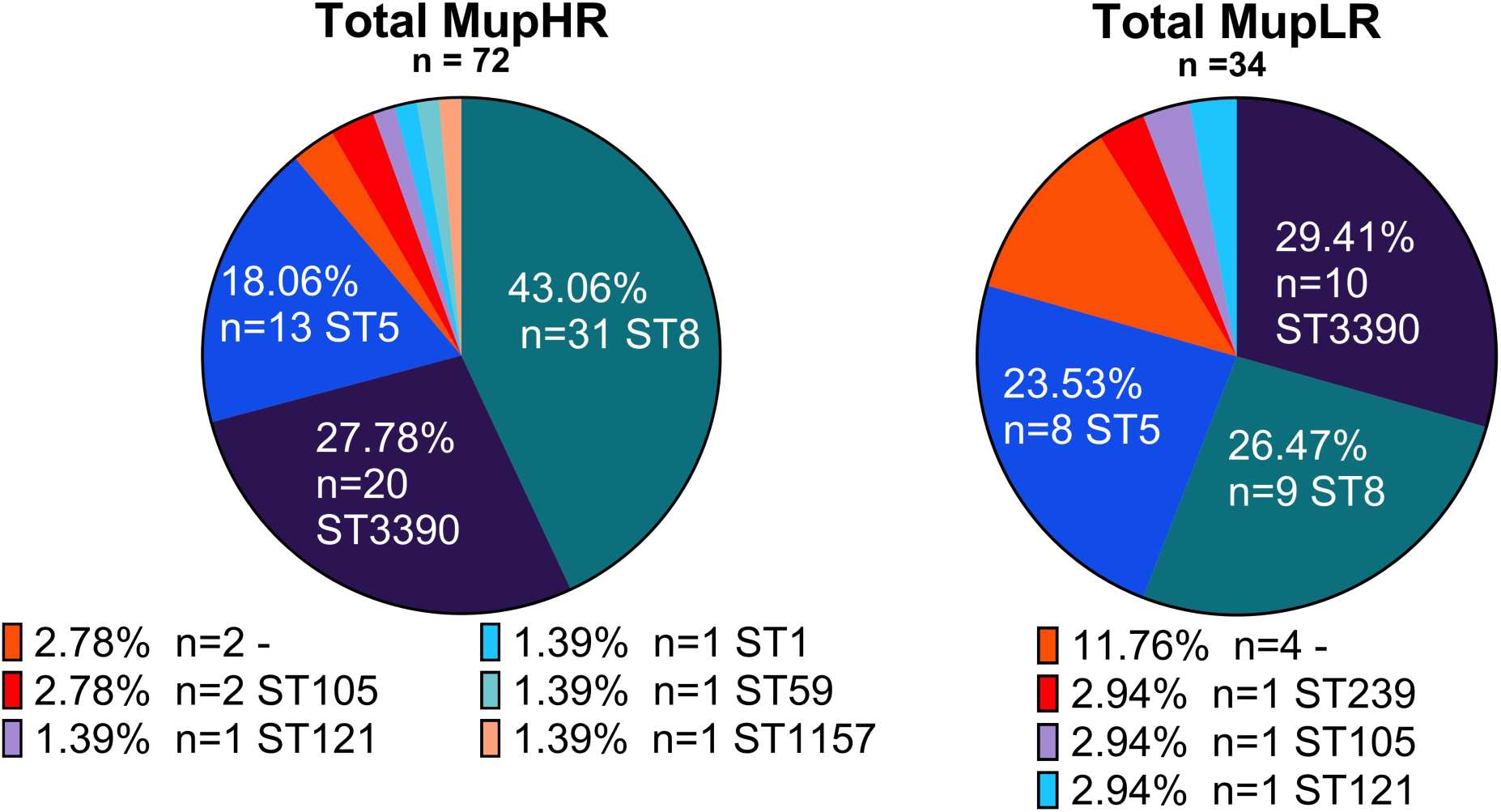
Mupirocin Resistance as a Function of Sequence Type. Shown are the sequence types for high-level mupirocin resistance (MupHR) and low/intermediate-level mupirocin resistance (MupLR) of the 106 MupR strains as determined by analysis of whole genome sequence. Mupirocin susceptibility was determined by Etests. Two MupHR and four MupLR strains were untypeable.

When we explored MupR as a function of population structure, the three most common STs ST8, ST5, and ST3390, made up the majority of the MupHR (88.9%) and MupLR (79.4%) strains (**Figure 3**). MupHR strains comprised eight different STs, with the highest proportion being ST8 (43.1%) with 31 strains (PreC = 19, 43.2% vs. PostC = 12, 42.9%), then ST3390 (27.8%) with 20 strains (PreC = 13, 29.5% vs. PostC = 7, 25%) and 13 ST5 strains (PreC = 8,18.2% vs. PostC = 17.9%). ST3390 made up the highest group of MupLR overall with 10 strains (PreC = 6, 35.3% vs. PostC = 4, 23.5%), followed by ST8, the majority of which are PostC (PreC = 2, 11.8% vs. PostC = 7, 41.2%), and then finally, ST5 with most being PreC (PreC = 7, 41.2% vs. PostC = 1, 5.9%). Other MRSA-MupHR were ST105, ST1, and ST59. Of the three MSSA-MupR, two were ST121 (one MupHR, one MupLR) and one ST1157 (Murph).

**Figure 3.**
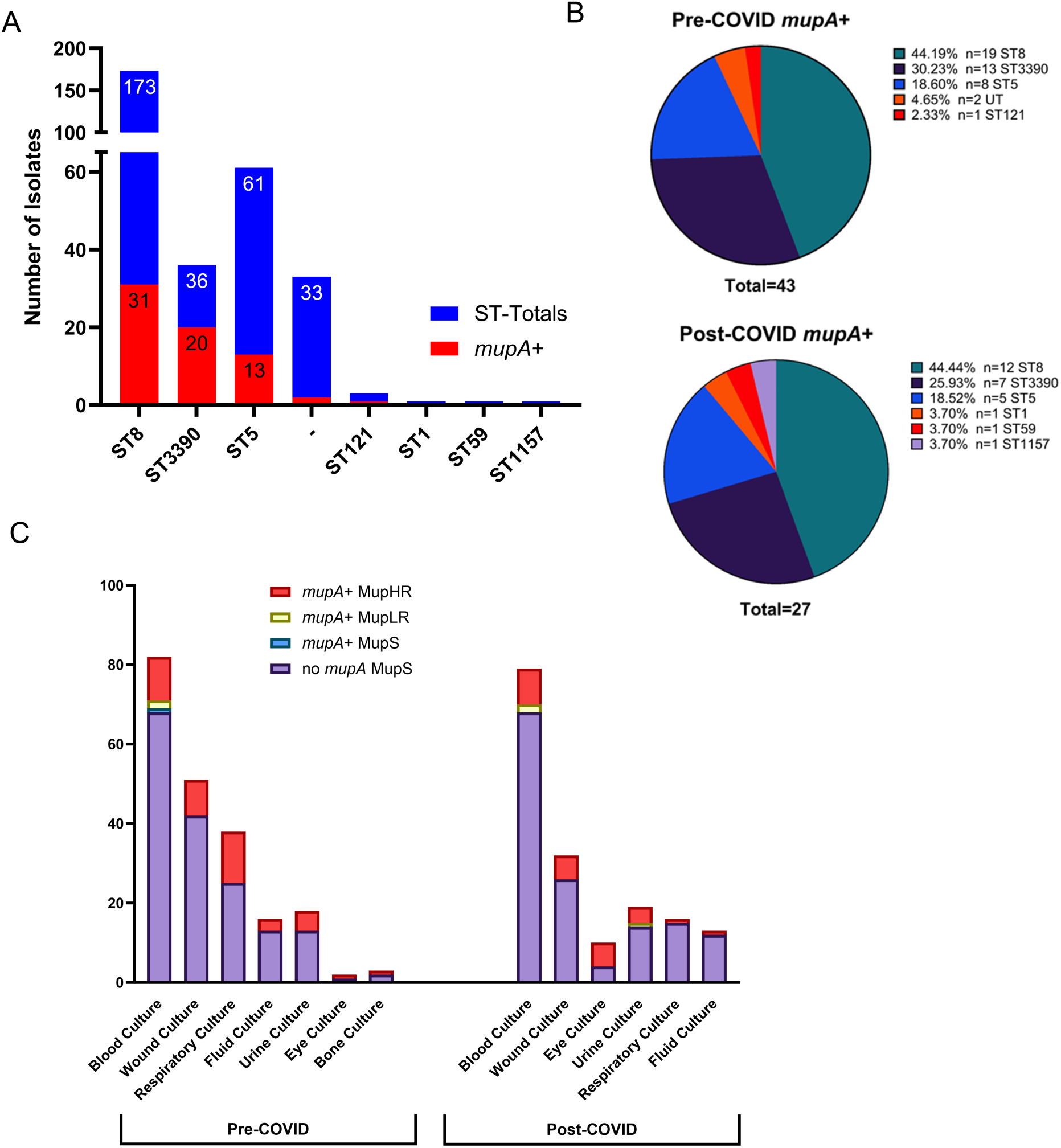
Distribution of *mupA* across Sequence Types and Correlation with Mupirocin Resistance. Summary of the distribution of *mupA^+^* isolates and ST across: (A) the entire 384 strain collection, (B) the Pre-COVID strains and the Post-COVID strains. Shown are the percentages and number of strains for each ST. C. The distribution of clinical strains with and without *mupA* and its phenotypic impact are shown. Most *mupA^+^* strains were MupHR as determined by eTest. Strains are arrayed by isolation site, isolation period, the presence-absence of *mupA*, and its phenotypic impact on mupirocin resistance/sensitivity. Strains with mutations in *ileS* that are phenotypically MupLR are not included in this depiction.

### High Level Mupirocin Resistant Strains Carry the *mupA* Gene

From our collection, 70 of 72 MupHR isolates contained the *mupA* gene, whilst no isolates were found to be *mupB* positive (**Figure 3A**). MupHR strains were most commonly classified in the top three STs within two CCs CC8 (ST8, n=31, 44.3%) and CC5 (ST5, n=13, 18.6% and ST3390, n=20, 28.6%). There was an overall decrease of *mupA* carriage over time (20.4% PreC vs. 15.6% PostC) that was not statistically significant (p=0.228). Within the pre-COVID strain collection (**Figure 3B**), we see a higher number of *mupA^+^* isolates (43 strains) from a less diverse mix of the population (four STs and two untypeable strains in total), whereas in the Post-COVID collection (**Figure 3B**) we see 27 strains distributed across six STs. The percentage of *mupA*^+^ ST8 (PreC = 19, 44.2% vs. PostC = 12, 44.4%) and ST5 isolates (PreC = 8, 18.6% vs. PostC = 5, 18.5%) remained largely stable, however there was a slight decrease in the percentage of ST3390 isolates (PreC = 13, 30.2% vs. PostC = 7, 25.9%, p-value = 0.264). The remaining *mupA*^+^ isolates made up a relatively small proportion of each population (PreC = 3, 6.9% vs. PostC = 3, 11.8%) and did not have any ST in common. The rates of *mupA* prevalence across the Pre and Post pandemic era were not found to be significantly different.

In general, the genotype and antibiotic susceptibility phenotype of our *mupA*^+^ strains conformed, e.g. *mupA*^+^ strains were MupHR and *mupA*^−^ strains were MupS (**Figure 3C**). However, there were exceptions. Isolate TGH406, was *mupA*+ but MupS, and no variation found in *mupA* sequence to indicate it had become a pseudogene. We also detected a cluster of five *mupA*^+^ isolates that were phenotypically MupLR (**Figure 3C**). We know from the literature that this is likely due to *mupA* integration onto the chromosome [8]. There were six samples that were found to have mutations within *mupA*, leading to varying degrees of protein truncations. Two truncations were found that resulted in 957/1024 AA (n=2) and 997/1024 AA (n=1) variants that did not have an impact on the function of *mupA* resistance measured phenotypically. Conversely, three isolates had larger truncations, all of 137 AA, resulting in a shorter protein sequence of 887 AA. This variant completely eliminated MupA function, resulting in a MupS phenotype.

### Mutations in *IleS* Correlate with Low Level Mupirocin Resistance

Low level mupirocin resistance is associated with a mutation (V588F) in the native isoleucyl-tRNA synthetase, *ileS*. This mutation was identified in 27/34 MupLR isolates (PreC = 14, 6.6% vs. PostC = 13, 7.5%). Three strains had the V588F mutation in *ileS* and also carried *mupA* and were MupHR as a result. There was also a single MupLR isolate (TGH365) that had a novel *ileS* mutation (W436G) that to our knowledge has never been reported. We also had four MupLR samples that did not contain any mutations in the *ileS* sequence.

From a population perspective, there was a notable switch in strains bearing the V588F mutation Pre-COVID and Post-COVID. Pre-COVID, there are no ST8 strains with MupLR that bear *ileS* mutations; instead, the most prevalent groups are ST5 and ST3390, represented by six isolates, each making up 42.9% of the population (**Figure 4A)**. In the Post-COVID population, ST8 represents 46.2% of the observed MupLR population, driven by the acquisition of the *ileS* (V588F) mutation (**Figure 4B)**. There are no ST8 samples with this mutation prior to this shift in the Pre-COVID population. Afterwards, Post-COVID ST8 made up 61.5% of all the strains with the (V588F) mutation. The discrepancy between the percentage of Post-COVID MupLR and Post-COVID (V588F) mutation ST8 strains is due to two ST8 strains in which the chromosomal mutation effect is masked by the presence of *mupA*, resulting in a MupHR phenotype. Thus, we observed CA-MRSA strains acquiring mutations in *ileS* and the *mupA* plasmid associated with MupHR.

**Figure 4.**
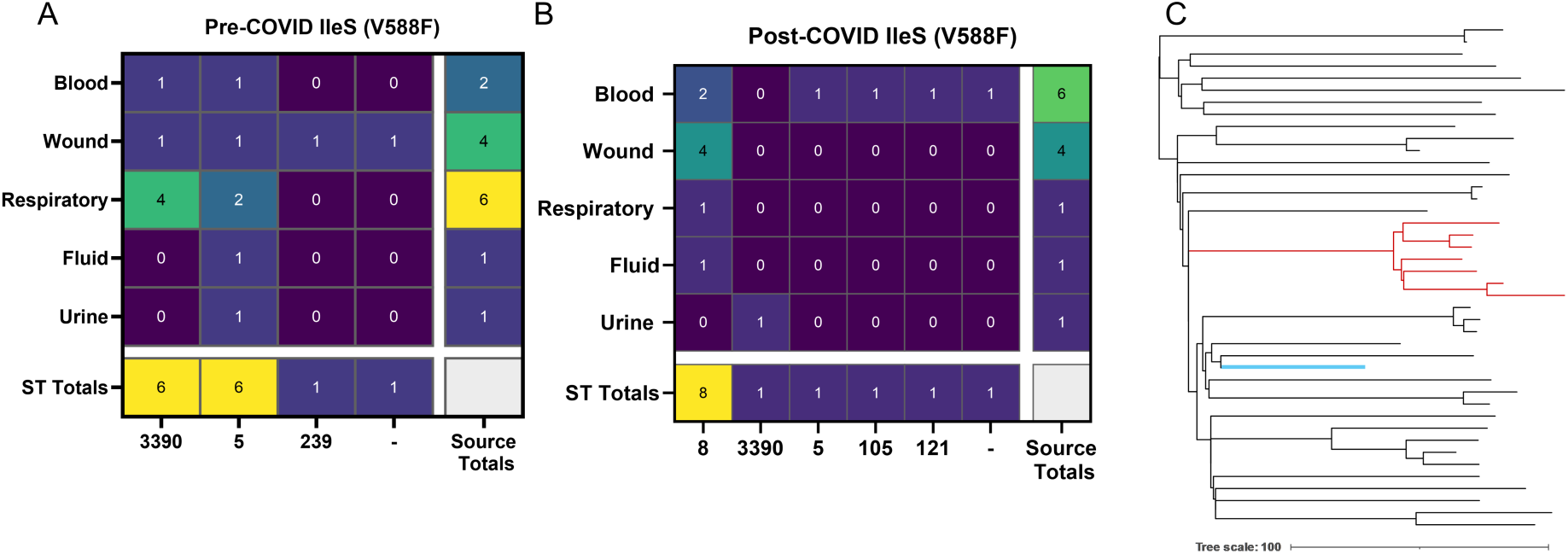
Correlation of MupLR and *ileS* (V588F) as a Function of Sequence Type and Isolation Site. Summary of the distribution of *ileS* (V588F) positive isolates **(A)** Pre-COVID and **(B)** Post-COVID. **(C)** Shown is a recombination masked core SNP alignment maximum-likelihood tree of Post-COVID USA300 isolates (ST8). The clade of 7 mupirocin resistant USA300 isolates that possess the R112Q *IleS* mutation are designated in red. The reference USA300 strain FPR_3757 used for SNP alignment is shown in blue.

In addition, we identified another conserved novel MupLR *ileS* mutation (R112Q) that has never been reported in the literature. This only appears within 7 of the 41 PostC USA300 strains, indicating that this cluster may represents an outbreak within the ST8 clone (**Figure 4C**). Over time the ST3390 population showed an increase in the presence of MupHR (+25.5% PostC), with decrease in samples with *ileS* mutations (-16.7% PostC), suggesting evolution to a more highly resistant population.

## Discussion

Prior studies have shown that the rate of MupR varies by geographic region and clinical characteristics of the patient populations sampled. The increasing use of decolonization protocols designed to reduce the incidence of MRSA transmission creates selection pressure for emergence of MupR. MupR has been reported in both MRSA and MSSA strains [9] [10], and genome sequencing has implicated transmission of strains within healthcare settings [9] [10]. Amongst our strain collection, prevalence of MupR was higher in MRSA isolates than MSSA isolates, as was also reported in strains from Stamford, CT [9]. For MRSA MupR strains, nasal povidone-iodine protocols for decolonization should be effective [29], although recent clinical trials have shown that nasal povidone iodine is inferior to nasal mupirocin [30]. Thus, optimal management of MRSA colonization is not clear and will be influenced by local molecular epidemiology of *S. aureus* and the rate of emergence of MupR.

The mechanism of MupHR is transmission of plasmid mediated *mupA* whilst MupLR is associated with accumulation of chromosomal point mutations. Amongst the risk factors reported for MupR are 1) prior exposure to mupirocin in the past year 2) *Pseudomonas aeruginosa* infection in the year prior to admission and 3) cefepime use in the year prior to culture [31]. Ceasing mupirocin exposure is reported to result in less MupR [8], although we did not see significant differences between rates of MupR Pre-COVID and Post-COVID. The Pre-COVID strain collection was less diverse than the Post-COVID collection so balancing selective pressures may be present. A limitation of our retrospective study is that we did not have a method to determine mupirocin exposure outside the inpatient setting. Other selective pressures that may be operative include exposure to other antibiotics that favor maintenance of the resistance plasmid. Interestingly, many of the isolates that carry *mupA* (58 out of 70), were also found to have a gene encoding for resistance to aminoglycoside antibiotics, ANT(4’)-Ia. Therefore, changing our decolonization protocol to povidone iodine may have slowed emergence of MupR, but this trend may have been masked by other factors affecting the population structure of MRSA in our hospital system.

The isolates examined were convenience samples, and our laboratory archives a random subset of clinical isolates. We noted differences MupR in some of the subpopulations of isolates tested, including respiratory samples. This phenotypic change in the Non-Blood category isolates may be due to a population shift over time from being predominantly CC5 (HA-MRSA) to CC8 (CA-MRSA), however, we cannot rule out that these differences were due to our small sample size. A more in-depth characterization of samples from different clinical sites is needed to determine if the changes we observed represent changes in population structure of clinical isolates, outbreaks or selection of specific clones.

There were two isolates (Blood/Respiratory) from the same patient that had MupHR with no identifiable *mupA* or *mupB* present. Both samples are ST105 SCC*mec* type II *Spa*-type t002, a relative of the New York/Japan clone (USA100/ST5). European ST105-II-t002 has been linked to skin and soft tissue infections, while the more recent South American ST105-II-t002 has been heavily associated with blood stream infections [32, 33].

A major theme in *S. aureus* pathogenesis has been the emergence of waves of virulent clones carrying genetic determinants conferring resistance to antibiotics [34]. In the 1990’s strains were either HA-MRSA or CA-MRSA, but gradually virulent CA-MRSA strains such as USA300 became dominant [34]. We show that MupR in western Florida is associated with MRSA strains and common US MRSA types such as ST8 (USA300 and USA500) and ST5 (USA100) are highly represented within our clinical strains. Our collection also revealed the emergence of an unusual ST3390 group with unexpected high prevalence of MupR (83%). Our ST3390 strains show greater toxicity to neutrophils and are more virulent in mice than reference ST5 strain (Felton, *et al* submitted). The genomes of ST3390 strains show evidence of horizontal exchange of genetic traits that could promote maintenance in the population, increased virulence, and multi-drug resistance (Felton, *et al* submitted).

Decolonization protocols for *S. aureus* utilizing antibiotics and antiseptics such as mupirocin or povidone-iodine with chlorhexidine have shown to decrease surgical site infections and invasive infections, but only result in temporary decolonization. These agents could promote the selection of resistant organisms through selective pressure or broad disruption of the microbiome. The incidence of mupirocin resistance and the molecular epidemiology of resistant strains in local hospital populations is an important factor in selecting the appropriate agents used for decolonization.

## Abbreviations

MRSA: methicillin resistant Staphylococcus aureus
MSSA: methicillin sensitive Staphylococcus aureus
MupR: Mupirocin Resistant
CC: clonal complexes
ST: sequence type

## Acknowledgements

This study was supported by grants AI124458 and AI157506 (both to L.N.S.) from the National Institute of Allergy and Infectious Diseases and a USF Provost’s CREATE award (LNS and KK). We thank Emily Coughlin of the MCOM RISE program with advice about statistical analysis, the USF Genomics facility, including John Adams and Min Zhang, for expert advice and assistance, and Kartik Cherabuddi for review of the manuscript.

## Data availability

Whole genome sequence data for all 384 isolates can be accessed at SRA with accession numbers <u>PRJNA785927</u> (Run1 80 isolates) and <u>PRJNA1125331</u> (Run 2 and 3; 304 isolates). Sequenced strains are listed in Supplementary Table 1.

---

**Supplemental Table 2.**
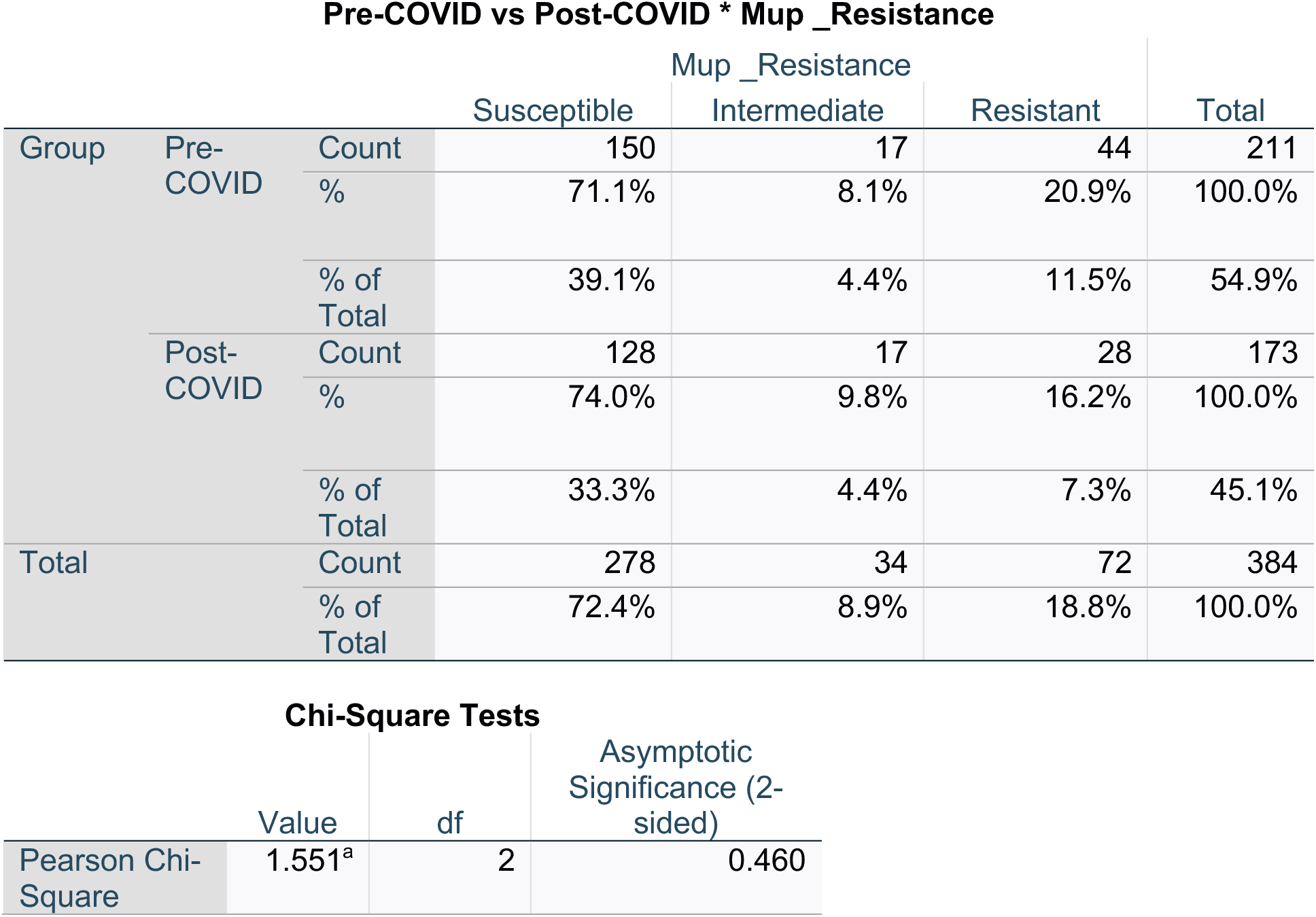
Mupirocin Resistance Comparisons.

**Supplemental Table 3.**
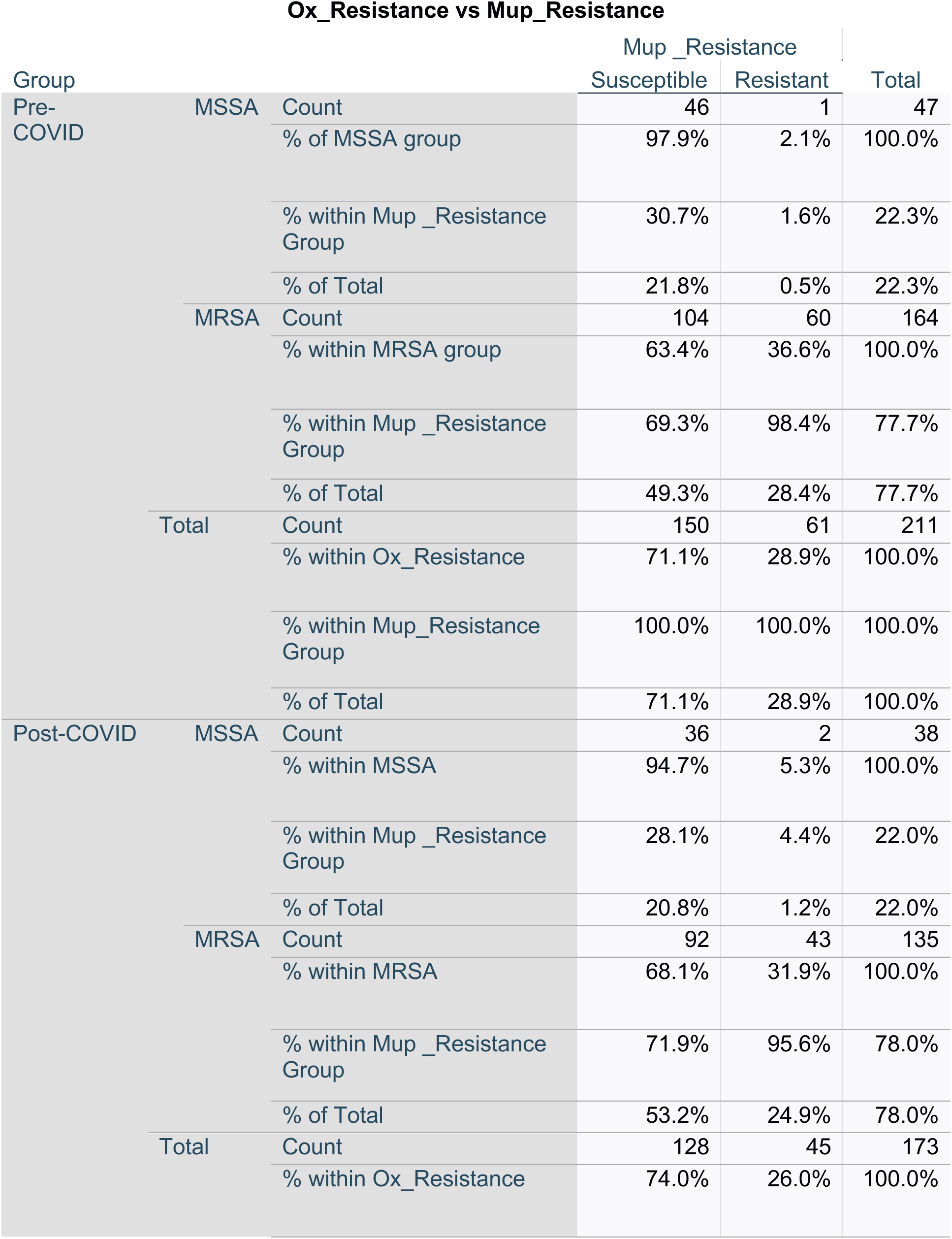
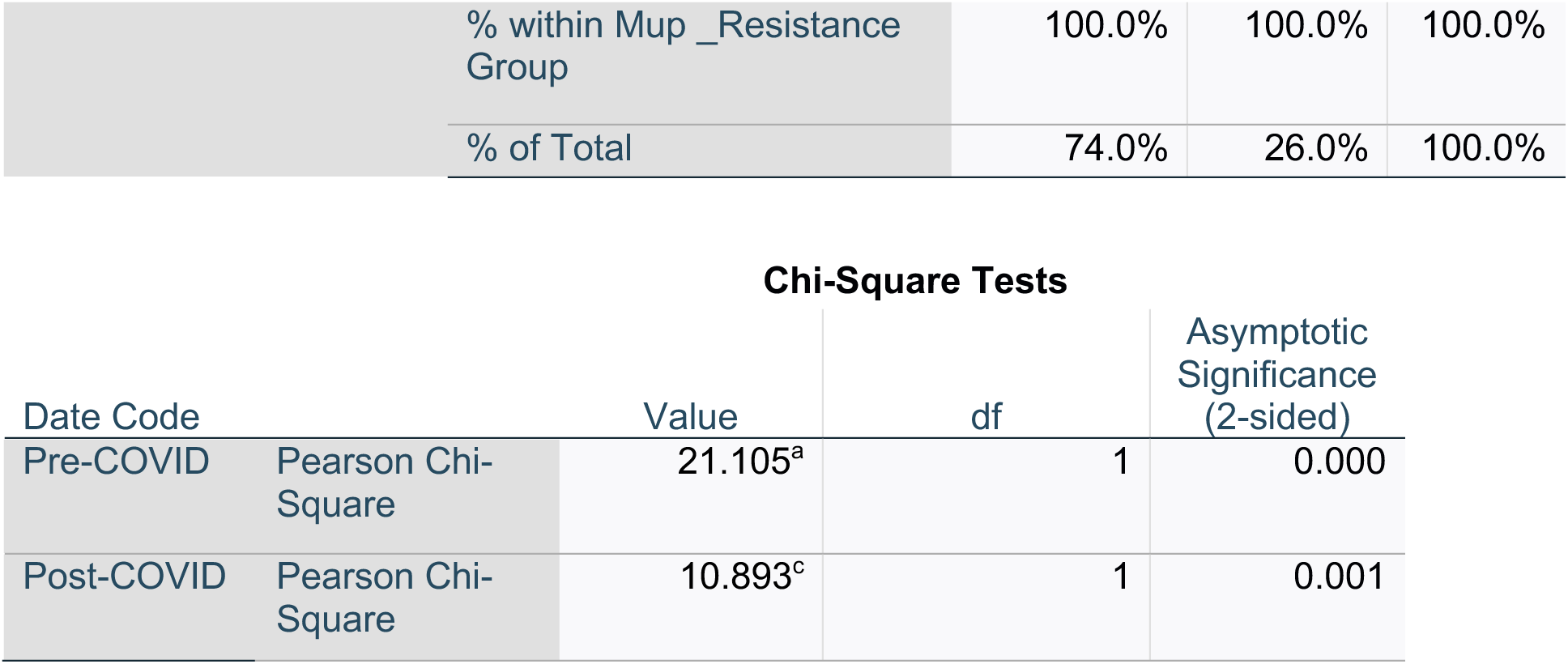
Ox\_Resistance vs Mup\_Resistance

| Group |  |  | Mup_Resistance |  | Total |
| --- | --- | --- | --- | --- | --- |
|  |  |  | Susceptible | Resistant |  |
| Pre-COVID | MSSA | Count | 46 | 1 | 47 |
|  |  | % of MSSA group | 97.9% | 2.1% | 100.0% |
|  |  | % within Mup_Resistance Group | 30.7% | 1.6% | 22.3% |
|  |  | % of Total | 21.8% | 0.5% | 22.3% |
|  | MRSA | Count | 104 | 60 | 164 |
|  |  | % within MRSA group | 63.4% | 36.6% | 100.0% |
|  |  | % within Mup_Resistance Group | 69.3% | 98.4% | 77.7% |
|  |  | % of Total | 49.3% | 28.4% | 77.7% |
|  | Total | Count | 150 | 61 | 211 |
|  |  | % within Ox_Resistance | 71.1% | 28.9% | 100.0% |
|  |  | % within Mup_Resistance Group | 100.0% | 100.0% | 100.0% |
|  |  | % of Total | 71.1% | 28.9% | 100.0% |
| Post-COVID | MSSA | Count | 36 | 2 | 38 |
|  |  | % within MSSA | 94.7% | 5.3% | 100.0% |
|  |  | % within Mup_Resistance Group | 28.1% | 4.4% | 22.0% |
|  |  | % of Total | 20.8% | 1.2% | 22.0% |
|  | MRSA | Count | 92 | 43 | 135 |
|  |  | % within MRSA | 68.1% | 31.9% | 100.0% |
|  |  | % within Mup_Resistance Group | 71.9% | 95.6% | 78.0% |
|  |  | % of Total | 53.2% | 24.9% | 78.0% |
|  | Total | Count | 128 | 45 | 173 |
|  |  | % within Ox_Resistance | 74.0% | 26.0% | 100.0% |

|  |  |  |  |  |
| --- | --- | --- | --- | --- |
|  | % within Mup _Resistance Group | 100.0% | 100.0% | 100.0% |
|  | % of Total | 74.0% | 26.0% | 100.0% |

**Chi-Square Tests**
| Date Code |  | Value | df | Asymptotic Significance (2-sided) |
| --- | --- | --- | --- | --- |
| Pre-COVID | Pearson Chi-Square | 21.105<sup>a</sup> | 1 | 0.000 |
| Post-COVID | Pearson Chi-Square | 10.893<sup>c</sup> | 1 | 0.001 |

**Supplemental Table 4.**
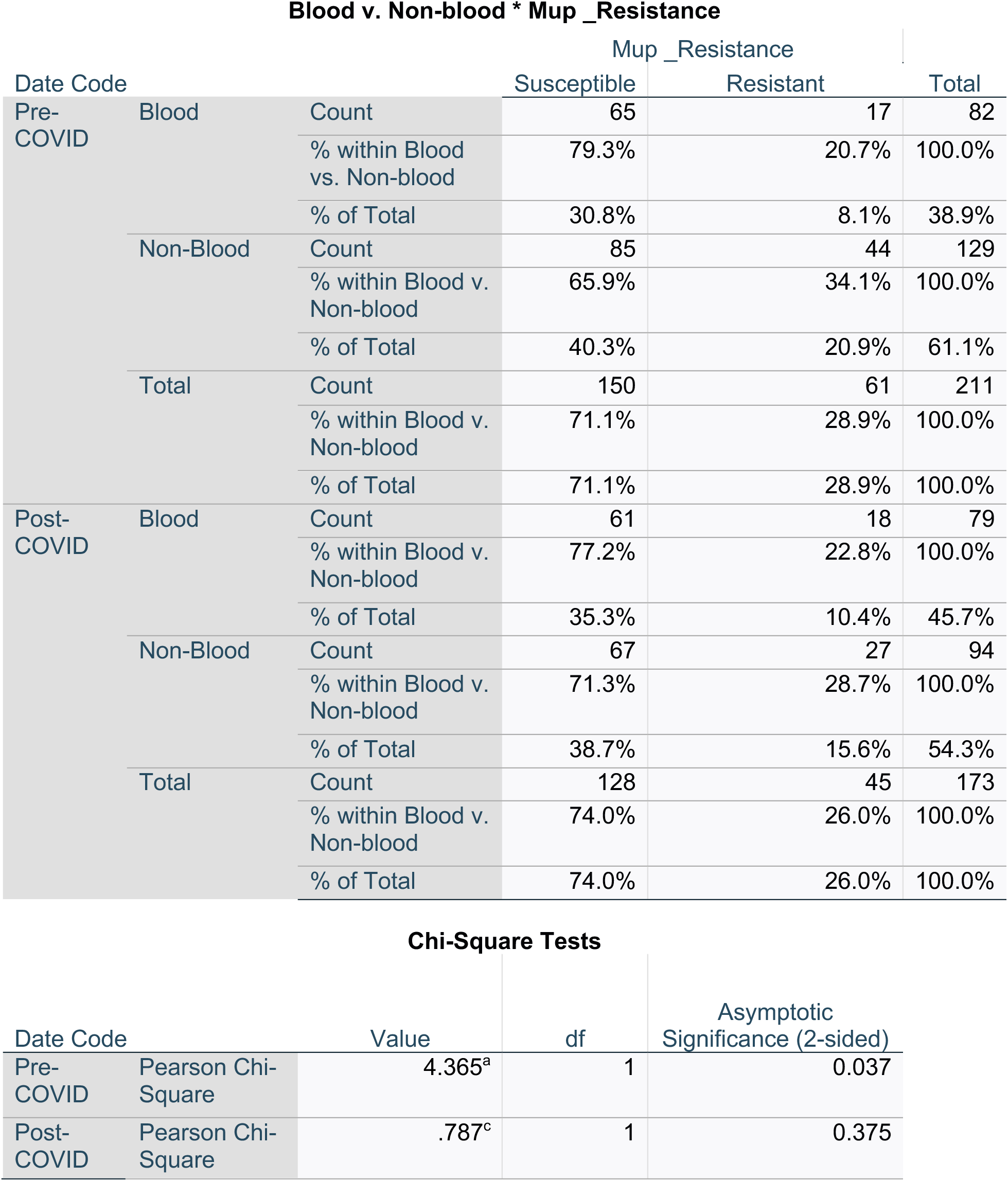

**Supplemental Table 5.**
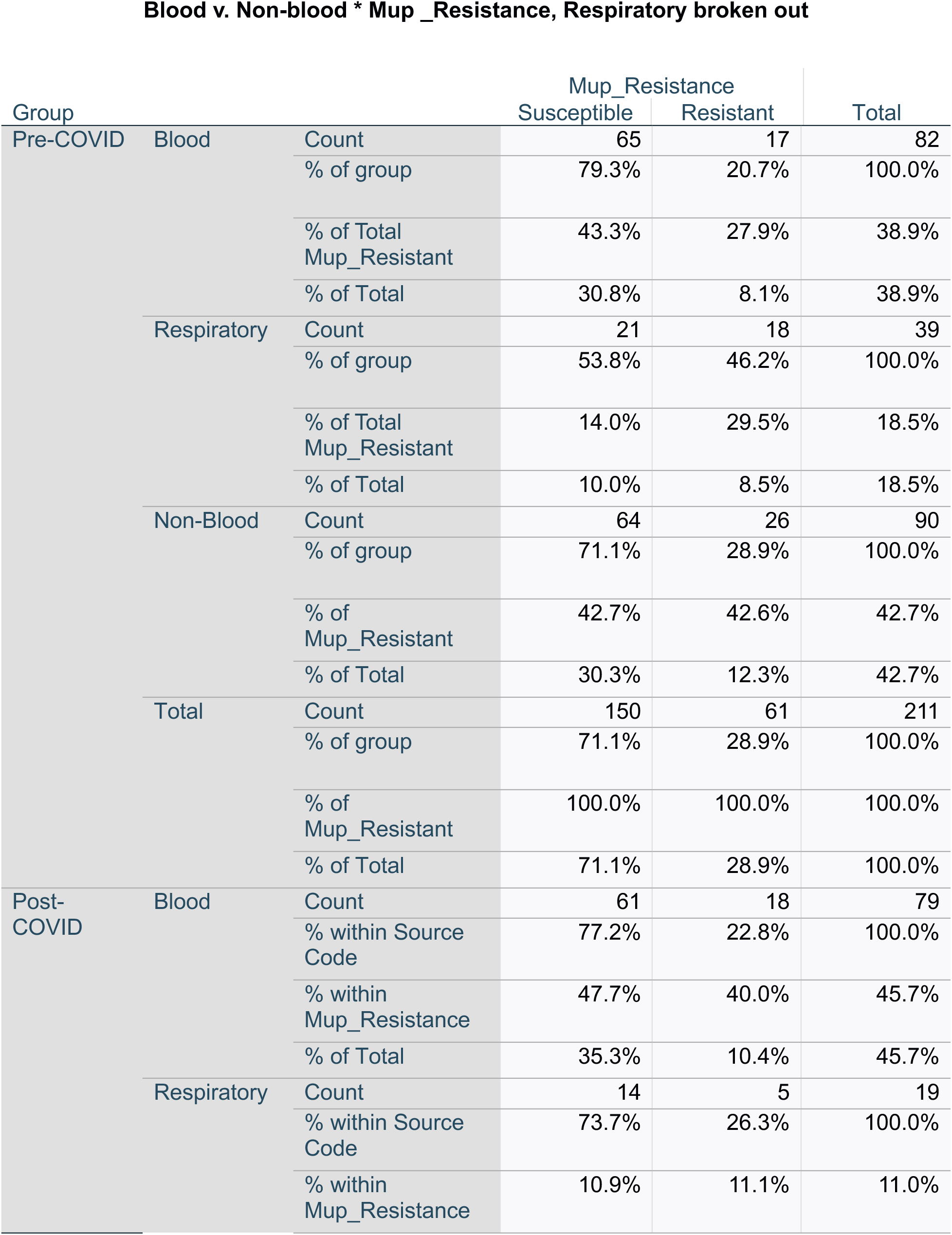
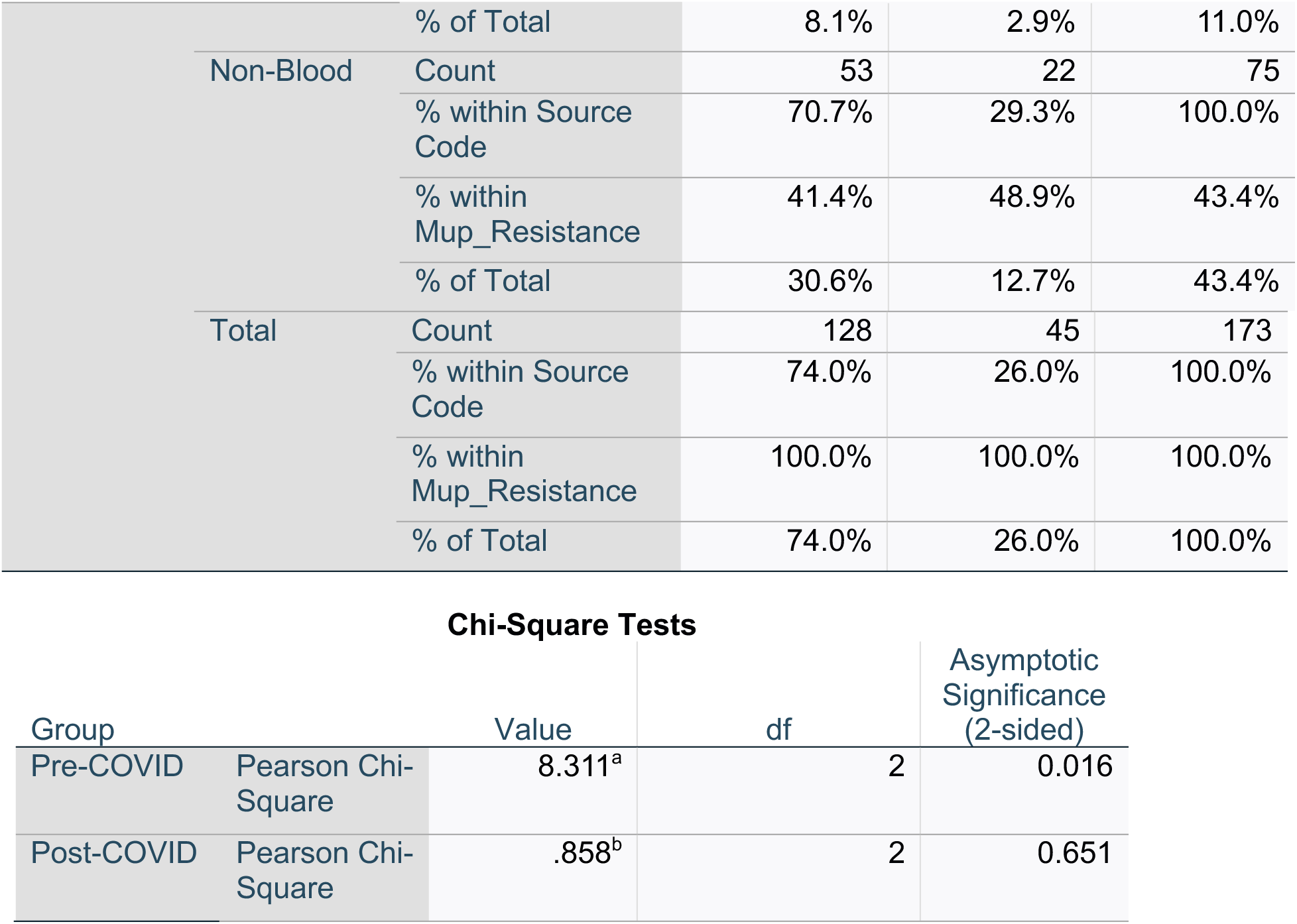

